# Efficacy and safety of a patch containing adipose-derived stem cells for skin wound healing – results form a comprehensive pre-clinical evaluation program

**DOI:** 10.1101/2022.03.09.483580

**Authors:** Nicolo C. Brembilla, Ali Modarressi, Dominik André-Lévigne, Estelle Brioudes, Florian Lanza, Hubert Vuagnat, Stéphane Durual, Laurine Marger, Wolf-Henning Boehncke, Karl-Heinz Krause, Olivier Preynat-Seauve

## Abstract

Mesenchymal stem cell-based therapies are emerging as innovative approaches to treat chronic wounds. A common administration route used in clinical trials consists of local injections leading to uncontrolled/sub-optimal delivery. This study reports a comprehensive pre-clinical evaluation program on the mechanism of action, efficacy and safety of an easy-to-use patch that concentrates Adipose-derived Stem Cells (ASCs) in a clinical-grade sponge of porcine crosslinked-gelatin. ASCs were prepared from the fat of ischemic patients. Transcriptome and proteome of ASC-patches and ASC monolayers were assessed by microarrays, bio-arrays and mass spectrometry. Tumorigenesis was investigated in immunosuppressed mice according to the European Pharmacopeia. Angiogenesis was assessed *in vivo* in the chick chorioallantoic membrane model. Efficacy of the ASC-patch was tested in a rat model of ischemic full-thickness skin defect. Cell stability was assessed by luminescence using ASC-patches generated from ASCs stably transduced with firefly luciferase. ASCs from ischemic patients upregulated the transcription of multiple genes involved in skin wound healing when cultured within the ASC-patch formulation. The patch was not only a concentrator, but also a reservoir of both ASC-derived regenerative factors and sponge-derived soluble fragments having healing capacity. The secretome of the ASC-patch promoted dermal fibroblast survival and epidermal epithelialization. No tumor formation was observed in immunodeficient Nude mice subcutaneously transplanted with the ASC-patch. Transplanted patches were early invaded by new vessels *in vivo*, and a marked angiogenesis was confirmed in two independent animal models. Finally, ASC-patches prepared from syngeneic rats promoted faster healing and re-vascularization of full-thickness skin defects in a rat animal model. Of note, ASCs were viable and locally stable for at least two weeks *in vivo*. We provide here compelling pre-clinical evidence that patches concentrating ASCs within crosslinked gelatin may represent a convenient and effective tool for the management of chronic wounds.

## INTRODUCTION

Cutaneous wound repair is a physiological process aimed at restoring a lost skin integrity thought a highly regulated and overlapping series of events, including hemostasis, inflammation, cell proliferation, angiogenesis and tissue remodeling. Chronic skin wounds arise from a dysfunction of this process, which may manifest in any of its constituent phases, and result in a skin sore which is unable to timely heal [1]. There is no established consensus for the duration of a wound that defines chronicity, although 4 weeks to 3 months is often suggested [2]. Several diseases can predispose to chronicity, thus chronic wounds are classified depending on underlying etiology, i.e. diabetic ulcers, venous or arterial wounds and pressure ulcers. Given the variety of causes, chronic wounds are frequent. Recent estimates indicate that they affect about 1-2% of the worldwide population, and up to 5% of subjects older than 65 years [3,4]. This number is still expected to grow due to an increasing prevalence of ulcer-causing diseases as well as the overall increased life expectancy. Chronic ulcers carry a heavy human and economic burden [5]. Patients, who suffer from severe pain, loss of function and mobility and destress, experience a dramatic impairment of their lives. Chronic infections and amputations are common complications.

The treatment landscape for the management of chronic wounds is particularly vast and spans from dressings, debridement, negative pressure to advanced skin replacement technologies [6,7]. While the efficacy of each of these therapies has been shown in specific settings, a unique gold-standard in chronic wound management is lacking. Even in the presence of guidelines, several ulcers are not efficiently managed with current therapies. The development of more effective approaches is imperative to diminish the current human and economic burden that chronic wounds impose to the society and the healthcare system.

The last decade has witnessed important advances in chronic wound treatment solutions. In this respect, cell therapy based on Mesenchymal Stromal Cells (MSC) has emerged as particularly promising [8]. MSC are multipotent stem cells that can differentiate into mesenchymal tissue lineages (including osteoblasts, chondrocytes, myocytes, and adipocytes). While MSC are found in many tissues, they are mostly enriched in the bone marrow and the adipose tissue. Several in vitro and pre-clinical animal studies have demonstrated that MSC exert beneficial effects in all phases of wound healing (reviewed in [9]). MSC promote hemostasis [10]; suppress excessive inflammatory responses through immunomodulation and initiate the switch of macrophages toward a pro-healing M2 phenotype [11-13]; increase survival, migration and proliferation of fibroblasts allowing ECM remodeling through secretion of collagens and Matrix Metallo-Proteinases (MMP) [14,15]; produce anti-fibrotic factors [16]; and promote neovascularization and vasoprotection [17]. Therapeutic effects of MSC were shown to depend mainly on paracrine mechanisms, via secretion of cytokines, chemokines and growth factors, and production of extracellular vesicles [18,19].

Early clinical trials have also confirmed that both bone marrow- and adipose tissue-derived MSC have a positive effect on chronic wound healing with a good safety profile, independently of the underlying disease causing the condition [20-22]. The promise of MSC therapy remains high, although some obstacles could compromise its large-scale application. Cell sourcing, delivery method and MSC stability *in vivo* are major challenges. While adipose tissue is emerging as the most convenient source thanks to its easy access [23], other challenges have yet to be solved. The most common delivery modality used in on-going trials relies on multiple intramuscular, intra-wound or peri-wound injections [9,21]. This administration route is however invasive and associated with intense pain, particularly when the injection is applied close to the lesion. Moreover, systemically administered MSC were shown to be initially entrapped within the lungs, before reaching the wound [24]. Only a minority of MSCs reaches the wound site and directly contributes to the wound repair when a systemic route is used, while the possible systemic effects of MSC remain partially unknown and need further clarification. Studies using animal models have in addition highlighted that the *in vivo* stability of systemically injected MSC is relatively low [25,26], possibly limiting their efficacy.

In this study we generated patches that concentrates ASCs within a clinical-grade hemostatic surgical sponge composed of crosslinked porcine gelatin. This formulation aimed at concentrating locally the ASC regenerative secretome in the wound bed. Crosslinked gelatin, which is physically stable but also biodegradable and biocompatible, has the capacity to capture charged proteins such as those secreted by ASC [27-30]. In addition, gelatin has direct healing properties and is highly hydrophilic, therefore well suited to absorb wound exudate, making it an ideal substrate for chronic wound cell therapy applications. The ASC-patch studied here represents an easy and convenient formulation to limit the constraints of current injection-based protocol of mesenchymal cell therapy for chronic wounds. Our study validates this approach in pre-clinical settings and paves the way to its use in first-in-human trials.

## METHODS

### ASC culture and engineering of an ASC-enriched gelatin patch

Human ASC were prepared from the fat of patients affected by limb ischemia (Uckay, I., et al., Regenerative Secretoma of Adipose-Derived Stem Cells from Ischemic Patients. J Stem Cell Res Ther, 2019. 9(5)), and were cultured in Dulbecco’s Modified Eagle Medium DMEM (4,5 g/l glucose, L-Glutamine) supplemented with 10% of human platelet lysate (Stemulate, Cook Regentek), 1% penicillin and streptomycin (Thermofisher) at 37°C and under 5% CO2. Rat ASC were grown in the same medium than human ASC supplemented with 10% of fetal calf serum (Thermofisher). To manufacture the ASC-enriched gelatin patch, a piece of sterile absorbable gelatin sponge USP Gelfoam ((2×6cm)×7mm, Pfizer) was soaked in a suspension of ASC at a final density of 6000 cells/mm^3^. Upon complete absorption, the ASC-soaked sponge was cultured on a PET-membrane insert (Greiner bio-one) floating on ASC culture medium. Alternatively, ASC were incorporated on a gel containing 2.3 mg/ml of rat type I collagen. Gel polarization was achieved at room temperature (RT) upon neutralization of the acidic collagen solution with bicarbonate at a final pH=7. Collagen gels were cultured at the air-liquid interphase at 37°C in presence of ASC media. This study was conducted according to the approval by local ethical committee of the University Hospitals of Geneva, Switzerland (2020-01102 and NAC 14-183). Written informed consent was obtained from each individual

### Flow cytometry and multipotent differentiation of ASC

Cells were incubated with fluorochrome-labeled antibodies for 30 minutes at 4°C in binding buffer (BD Biosciences), prior to analysis using a BD Accuri™-B6 flow cytometer (BD Biosciences). Antibodies were as follows: i) for human cells: mouse IgG1 anti-CD44/CD73/CD90/CD45/CD105/CD14/HLA-DR (all from Abcam); ii) for rat cells: Armenian hamster anti-rat CD29-APC (clone HMb1-1, ThermoFischer), mouse anti-rat-CD31-PE (clone TLD-3A12, BD Pharmingen), mouse anti-rat CD45-BV421 (clone OX-1, BD Pharmingen) and mouse anti-rat CD90-BB515 (clone OX-7, BD Pharmingen). Analysis was performed on viable cells (negative for Draq7), upon exclusion of cell doublets. Cell purity was >98%.

The multipotent differentiation into adipocytes, osteocytes and chondrocytes was performed by using the Human Mesenchymal Stem Cell Functional Identification Kit (R&D Systems) according to the supplier’s instructions.

### Immunocytochemistry and immunofluorescence on tissue sections

ASC were cultured on glass coverslips prior to fixation with paraformaldehyde 0.5% for 15 minutes at RT. Cells were incubated overnight (o/n) at 4°C in PBS containing 0.3% Triton X-100 and 0.5% bovine serum-albumin with the following primary antibodies: mouse IgM anti-Stro-1 (Clone STRO1, Thermo scientific). Detection was achieved using an anti-mouse IgM-Alexa 555 antibody for one hour at +4°C. Cells were stained with DAPI 1μg/ml for 10 minutes prior to final washing and mounting.

For histological analyses, tissues were washed in PBS and fixed with a 4% paraformaldehyde solution for 20 minutes prior to dehydration and embedment in paraffin. Upon rehydration, slides were stained in PBS supplemented with bovine serum albumin 1%, Triton X-100 0.3% o/n at 4°C with a mouse IgG anti-CD31 (Abcam). Upon staining with anti-mouse IgG-Alexa 555 antibody, slides were counterstained with DAPI and mounted in FluorSave medium (Calbiochem). Hematoxylin & Eosin staining and Masson’s trichrome staining were performed according to the standard protocol.

Immunocytochemistry and immunofluorescence applied to osteocytes, chondrocytes and adipocytes derived from ASC was performed with the reagents of the human mesenchymal stem cell functional identification kit (RnDSystems)

### Cytokine measurements

Cytokines were measured in the supernatants from ASC cultures and ASC-enriched patches using the human cytokine base kit A (R&D Systems) combined with a magnetic Luminex assay (Bio-plex 200, Biorad) according to the manufacturer’s instructions.

### Microarrays

Isolation of total RNA was performed by using RNeasy kit from Qiagen according to the manufacturer’s instructions. RNA concentration was determined by a spectrometer (Thermo Scientific™ NanoDrop 2000) and RNA quality was verified by 2100 bioanalyzer (Agilent).

Human and rat microarray was performed with the Clariom™ S Assay’s for human and rat, respectively, (Thermofisher) using the Complete GeneChip^®^ Instrument System, Affymetrix. Hierarchical clustering and principal component analysis were computed using TAC4.0.1.36 software (Biosystems) using the pheatmap package (https://cran.r-project.org/web/packages/pheatmap/index.html) with default settings. Gene Set Enrichment Analysis (GSEA), desktop version 3.0 was used to analyze the pattern of differential gene expression between the ASC-patch and the monolayer condition. The Gene Ontology Biological Process (GOBP) gene set from the Molecular Signatures Database was used. Result of GSEA analysis were visualized for enrichment map using Cytoscape 3.8.2.

### Mass spectrometry

Cultured human ASC or ASC-enriched patches were incubated for 45 minutes with collagenase NB6 (Nordmark) at 0.3 U/ml, washed with a serum-free DMEM (Thermofisher) and cultured for 24h at 37°C in serum-free medium. Upon clarification of supernatants at 500 x g for 10min, proteins were precipitated, digested and peptides analyzed by nanoLC-MSMS using an easynLC1000 (Thermofisher) coupled with a Q Exactive HF mass spectrometer (Thermofisher). Database searches were performed with Mascot (Matrix Science) using the Human Reference Proteome database (Uniprot). Data were analyzed and validated with Scaffold (Proteome Software) with 1% of protein FDR and at least 2 unique peptide per protein with a 0.1% of peptide FDR.

### Functional tests on skin cells (fibroblasts and keratinocytes)

Human foreskin fibroblasts (provided by Neurix SA, Plan-les-Ouates) were cultured in 24-well culture plates in DMEM culture media (4,5 g/l glucose, L-Glutamine) supplemented with 10% fetal calf serum (Gibco) and 1% penicillin-streptomycin (Gibco). Upon washing to remove eventual cell debris, fibroblasts were cultured for 18 hours in serum-free media in the presence or absence of conditioned media from ASC cultures (serum-free medium pre-incubated with cultured ASC or ASC-enriched patches) at the concentration of 720 000 cells/ cm2. The number of fibroblasts was counted by trypan blue exclusion.

Primary human keratocytes were obtained from the skin of individuals undergoing abdominoplastic surgery as previously described [31]. Cells were seeded onto an ASC-enriched patch and initially cultured in a submerged manner in fully-supplemented CnT-Prime Medium (CELLnTEC, Bern, Switzerland). Medium was subsequently changed to CnT-Prime 3D Barrier Medium (CELLnTEC) for one day, before lifting the ASC-enriched patch to the air-liquid interface to induce cell differentiation for 10 days.

### Chick chorioallantoic membrane model

Fertilized chicken eggs were incubated at 37 °C and placed with the smaller convexity pointing upward from ED1 (Embryo Development day) to ED4. At ED4, a hole was drilled through the smaller convexity pointing of the shell. At ED 7, the eggs were opened with scissors through the hole and the inner membrane to create a round window with approximate 1cm diameter. The developing chorioallantoic membrane was then irritated through creation of a micro-hemorrhage. With ASC in suspension, a silicon ring with 4 mm inner diameter was placed on the site of the generated hemorrhage. The ASC-enriched patched, fibroblast-enriched patches or control empty patches were deposited directly on the site of the generated hemorrhage. After implantation the window in the eggshell was covered with a paraffin film and placed in the incubator at 37°C. The number of vessels connexions to the patch were counted manually under a binocular loop.

### Animal experiments

The local veterinary authority approved all procedures in accordance with Swiss guidelines (authorizations GE-83-19 and GE-97-19).

For stability/tumorigenicity assays, ASC-enriched patches or Hela cells (5.10^5^ cells) were subcutaneously transplanted in the right flank of Nu/Nu mice and followed for 12 weeks for tumor palpation and necropsy according to the guidelines of the European pharmacopeia (EMEA/149995/2008).

The model of ischemic wound in the rat was performed as previously described [25,32]. Briefly, Wistar female rats of 250-300 g were pre-anesthetized by inhalation of isoflurane 5%, and anesthetized at the dose of 2%. Hairs were removed from the inguinal region using a mechanical shaver. All surgical procedures were performed under an operating microscope. Through a longitudinal incision made in the upper part of the left thigh, the external iliac and femoral arteries were dissected free along their entire length, from the common iliac to the saphenous artery, and one cm-length artery was removed. Immediately after the arterial resection, a wound was created on the dorsal aspect of the feet in all animals by removing a full-thickness skin area of 1.2 × 0.8 cm. Treatments were applied a day after the surgery. To maintain the patches on the wound, a gutter of perforated silicone interface (Mepitel, Molnlycke) was covered with a thin sheet of polyurethane (Opsite, SmithNephew) and sutured around the wound. The patches were removed at day 7 and the wound covered with polyurethane until full recovery. Daily macroscopic evaluation of the limbs and feet as well as wounds planimetry were performed until complete wound healing. All animals that deviated from the protocol were excluded from the analysis.

## RESULTS

### Generation of an ASC-enriched gelatin patch (ASC-patch)

While holding great promise for the treatment of chronic ulcers, adipose-derived MSC therapy (i.e ASC-therapy) currently relies in sub-optimal and inconvenient injection delivery methods. To develop an easy and efficient way to deliver ASC to the wound, we generated an ASC-enriched patch containing crosslinked gelatin. Human ASC, prepared from fat tissue of patients presenting limb ischemia [33], were seeded into a clinical-grade (i.e Good-Manufacturing Practice, GMP-grade) porcine gelatin sponge and cultured in air/liquid interface conditions (Supplementary Figure 1A). After culture we obtained a stable and easy-handling cellular dressing (Figure 1A, left), characterized histologically by a dense cellular tissue (purple) interspersed between gelatin trabeculae (dark pink) (Figure 1A, middle panel). A green halo around gelatin trabeculae suggested collagen dissolution as assessed by Masson’s trichrome staining. ASC that clustered in pores started to organize into a compact tissue composed of collagen fibers from their own secretion (figure 1A, right panel). This tissue was the result of ASC-dependent secretion of extracellular matrix and contraction of the gelatin sponge, as only isolated cells in a rarefied gelatin mesh were observed at the beginning of the culture (Supplementary Figure 1B). A tissue characterized by lower cellular density was obtained upon parallel culture of ASCs within a collagen matrix, confirming the superiority of the gelatin/sponge formulations for ASC concentration (Supplementary Figure 1C).

**Figure 1.**
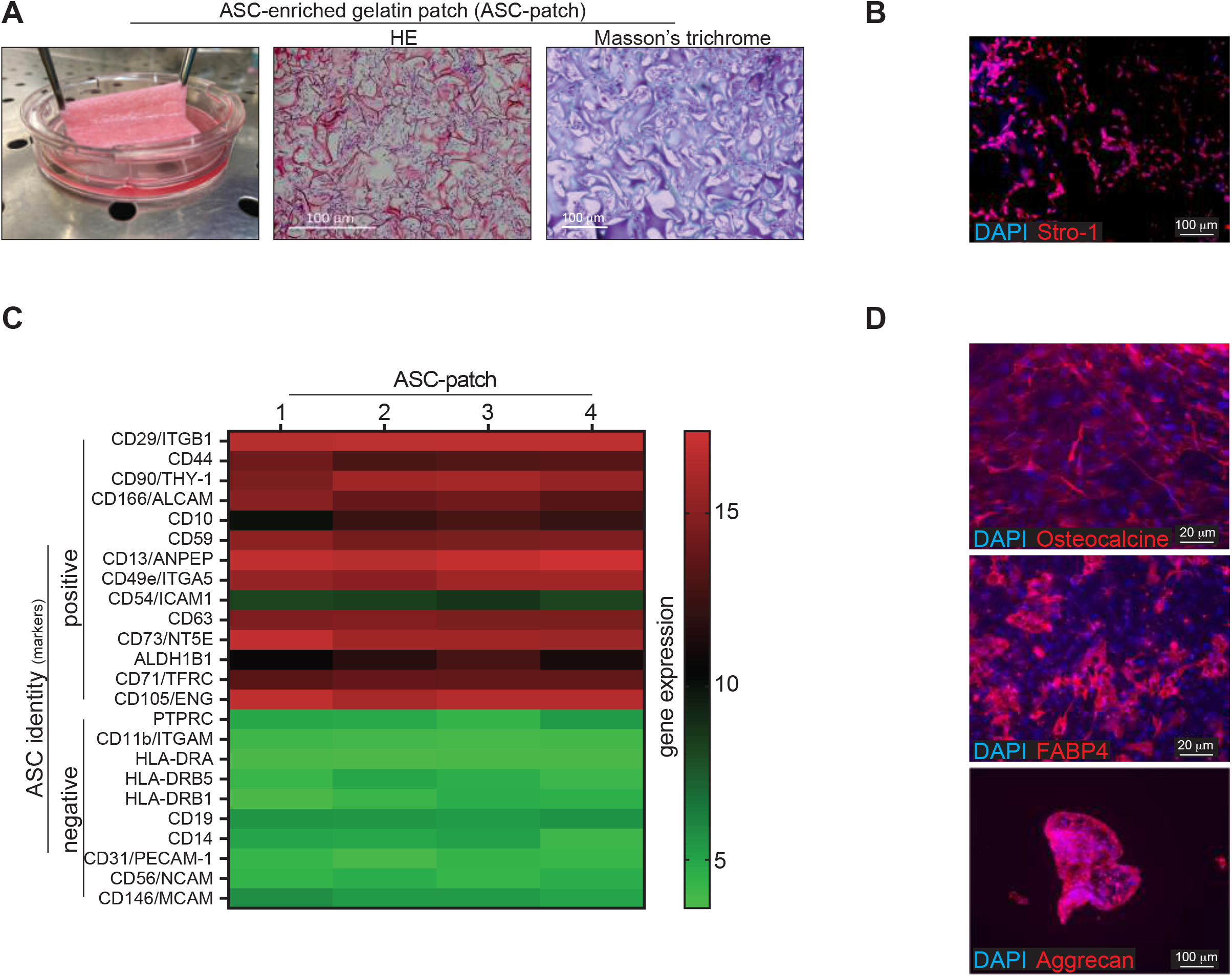
ASC cultured within a gelatin sponge generates a stable and easy-handling cellular patch (ASC-patch). A) Macroscopic image of an ASC-patch (left). Hemalum/eosin coloration (middle) and Masson-trichrome staining (right) of an ASC-patch are shown. B) Immunofluorescence staining of an ASC-patch section for the stromal marker Stro-1. C) Full gene expression array of ASC-patches generated from 4 independent ischemic patients. Commonly used positive markers for ASC identification are shown. D) Differentiated towards osteoblasts, adipocytes or chondrocytes of ASCs extracted from the ASC-patches. Osteocalcin (osteoblasts), FABP4 (adipocytes) and aggrecan (chondrocytes) staining are shown.

Of note, after one week of culture within the patch, ASC retained their stromal identity, as shown by sustained and widespread expression of the stromal marker Stro-1 (Figure 1B). Furthermore, cells obtained upon enzymatic dissociation of the ASC-enriched patch had a transcript profile compatible with undifferentiated ASC as defined in accordance to the current guidelines [34] (Figure 1C). Confirming the ASC identity, cells extracted from the ASC-enriched patch retained multipotent capacity, being able to differentiate towards osteoblasts (expressing osteocalcin), adipocytes (FAB4) and chondrocytes (aggrecan) in appropriate medium conditions (Figure 1D).

Together, we show here that ASC can efficiently be included and grown within a GMP-grade porcine gelatin support to generate an undifferentiated ASC-enriched compact tissue which has physical properties compatible with its use as a cellular patch.

### The ASC-enriched patch results in an enhanced regenerative transcriptome due to the interaction between ASC and gelatin

We next investigated whether ASCs have modified their gene expression capabilities as a result of their interaction with gelatin within the 3D patch culture conditions. To this aim, the entire transcriptome of cells extracted from three independent ASC-enriched patches was compared with ASC from the same batch but grown in monolayers. This latter condition was performed in line with the standard protocol used to produce ASC for injection based MSC therapy [35]. Row data are provided in Supplementary table 1. Hierarchical clustering and principal-component analysis (PCA) of microarray data revealed that ASC partially modified their global transcriptome when cultured within the ASC-enriched patch (Supplementary Figure 2A and B, respectively). ASC grown in monolayer for 7 days maintained a transcriptome profile similar to cells prior to culture (freshly isolated ASC vs monolayer (d7), Supplementary Figure 2B). Whether the condition used, ASCs did not differentiate in fibroblasts nor embryoid bodies, as such cells expressed characteristically different transcriptomic profiles (Supplementary Figure 2B).

To explore the characteristics of the genes expressed in ASC-patches compared to ASCs grown in monolayer as detected by microarray, we performed a threshold-free gene set enrichment analysis (GSEA). The 50 most highly differentially expressed genes (top 25 upregulations and top 25 downregulations) are shown in Figure 2A. Of note, pathways that were significantly most up-regulated were linked to proliferation, immune functions, morphogenesis and vascular growth, all being critical component of the healing process (Figure 2B). Down-regulation showed mainly a cluster linked to the regulation of DNA replication and one to mitochondria functions. Smaller clusters are shown in Supplementary Figure 2C.

**Figure 2.**
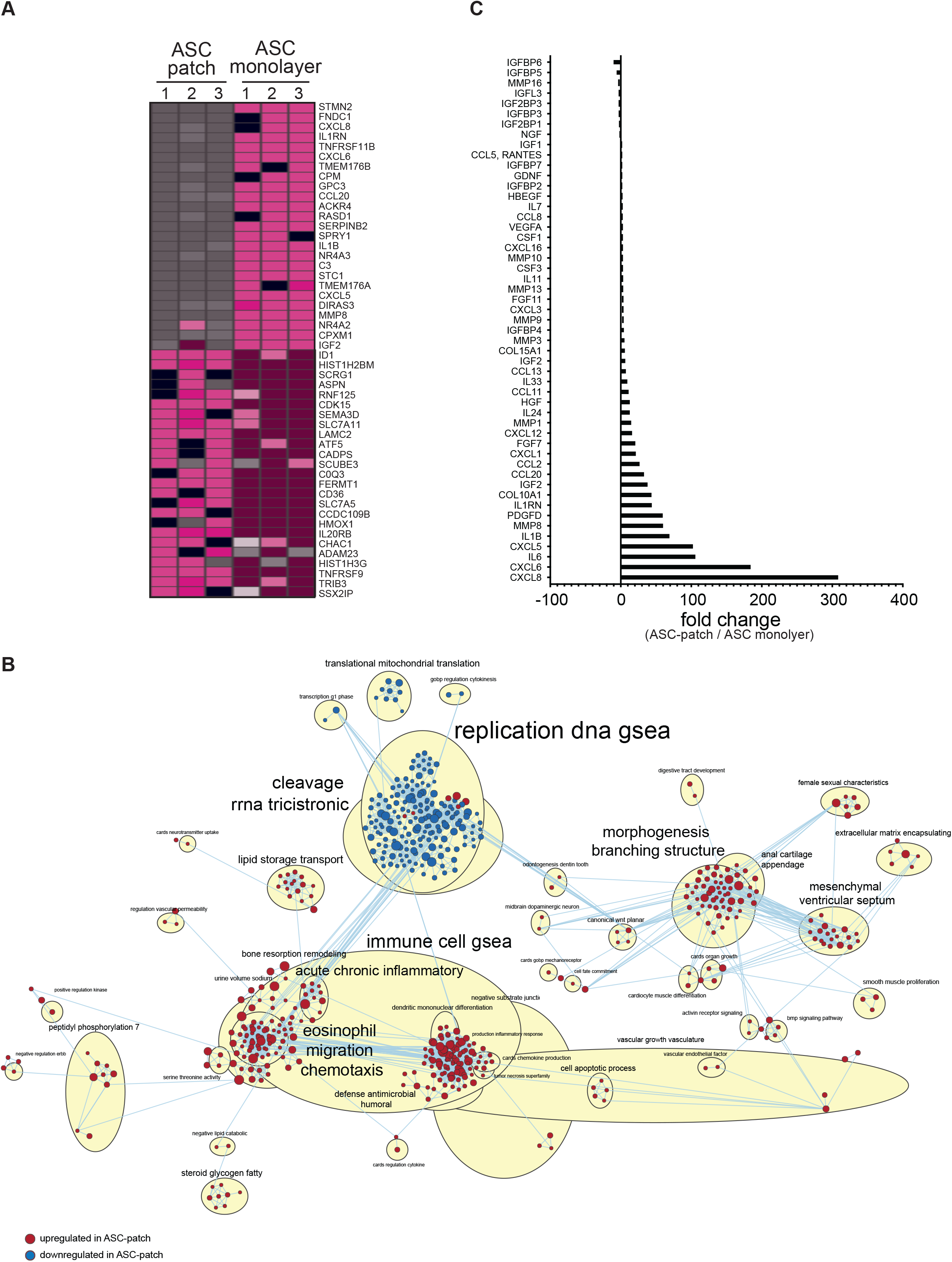
ASCs within the ASC-patch are characterized by an enhanced regenerative transcriptome. The transcriptome of ASC-patches generated from 3 ischemic patients were assessed by microarray and compared to ASC from the same donors grown in monolayer. A) Heatmap of the top 25 genes with increased expression and the top 25 with decreased expression in the ASC-patch compared with ASC monolayer condition, ranked by Gene Set Enrichment Analysis (GSEA). B) Enrichment pathways determined using the Gene Ontology Biological Process (GOBP) gene set from the Molecular Signature Database are visualized. C) Fold change analysis on 215 selected genes involved in wound healing. The 51 statistically significant regulations are shown (fold change >2 (up) and <2(down) and p<0.05, non-parametric Mann Whitney t test).

We next restricted and focus the analysis on 191 gene transcripts corresponding to the most implicated factors in the wound regeneration process (supplementary Table 2). Fifty-one out of the 191 analyzed transcript were significantly regulated in the ASC-patches (fold change >2 and p<0.05) compared with ASCs in monolayer: 30 strongly up-regulated (4x to 60x), 13 minimally up-modulated (<4x), 5 minimally down-modulated (<4x) and 3 modestly down-regulated (4x to 10x) (figure 2C). The strongest up-regulated transcripts observed were CXCL8 (IL-8) (potent angiogenic, chemotactic and inflammatory cytokine), CXCL6 (angiogenic, chemotactic, anti-microbial), IL-6 (angiogenic, pro-inflammatory), CXCL5 (angiogenic, matrix remodeling factor, pro-inflammatory). Other angiogenic factors were up-regulated (VEGF, ANGPTL2, ADM), as well as matrix remodeling factors (MMP16), collagens, cell growth factors (HGF) and several chemokines. Some IGF-Binding Proteins (IGFBP) were slightly down-regulated. Of note, the Keratinocytes Growth factor (KGF or FGF-7), promoting epithelialization during skin wound healing, was upregulated.

Together, these observations show that interaction of ASCs with gelatin within the ASC-patch results in a general enhancement of their regenerative transcriptome.

### The secreted proteome of ASC is absorbed by the gelatin sponge and concentrated within the ASC-patch

The composition of the proteome secreted in supernatants from the ASC-patch was profiled by mass spectrometry and compared to that released by the same ASC grown in monolayer, or the empty sponge. Hierarchical clustering analysis allows to clearly distinguish the three conditions (Figure 3A). The number of total proteins found in the supernatants of the ASC-patches was lower than the simple addition of the proteins found in the supernatants of the ASC-grown in monolayers plus those released by the empty sponge (Figure 3B). This observation may be indicative of protein adsorption within the patch. In line with this evidence, the protein diversity was slightly higher in supernatants from ASC grown in monolayer if compared to supernatants from ASC-patches (Figure 3A and 3C). More precisely, 3.1% of proteins were down-regulated and 2.7 up-regulated in the supernatants of ASC-patches as compared to the ASC monolayer’ supernatants (The complete list of proteins found in each condition can be found in supplementary table 3). Precise quantification by bio-arrays of cytokines, growth and angiogenic factors which have a key role in wound healing showed that these proteins were generally concentrated in the supernatant from ASC-patch (supplementary figure 3). Analysis of the supernatants from empty patches (no ASCs) additionally revealed that the gelatin slowly dissolves, releasing 49 proteins on average, most of which were collagens and keratins (supplementary figure 4). Of note, these proteins were also found in the supernatants from the ASC-patch (data not shown).

**Figure 3.**
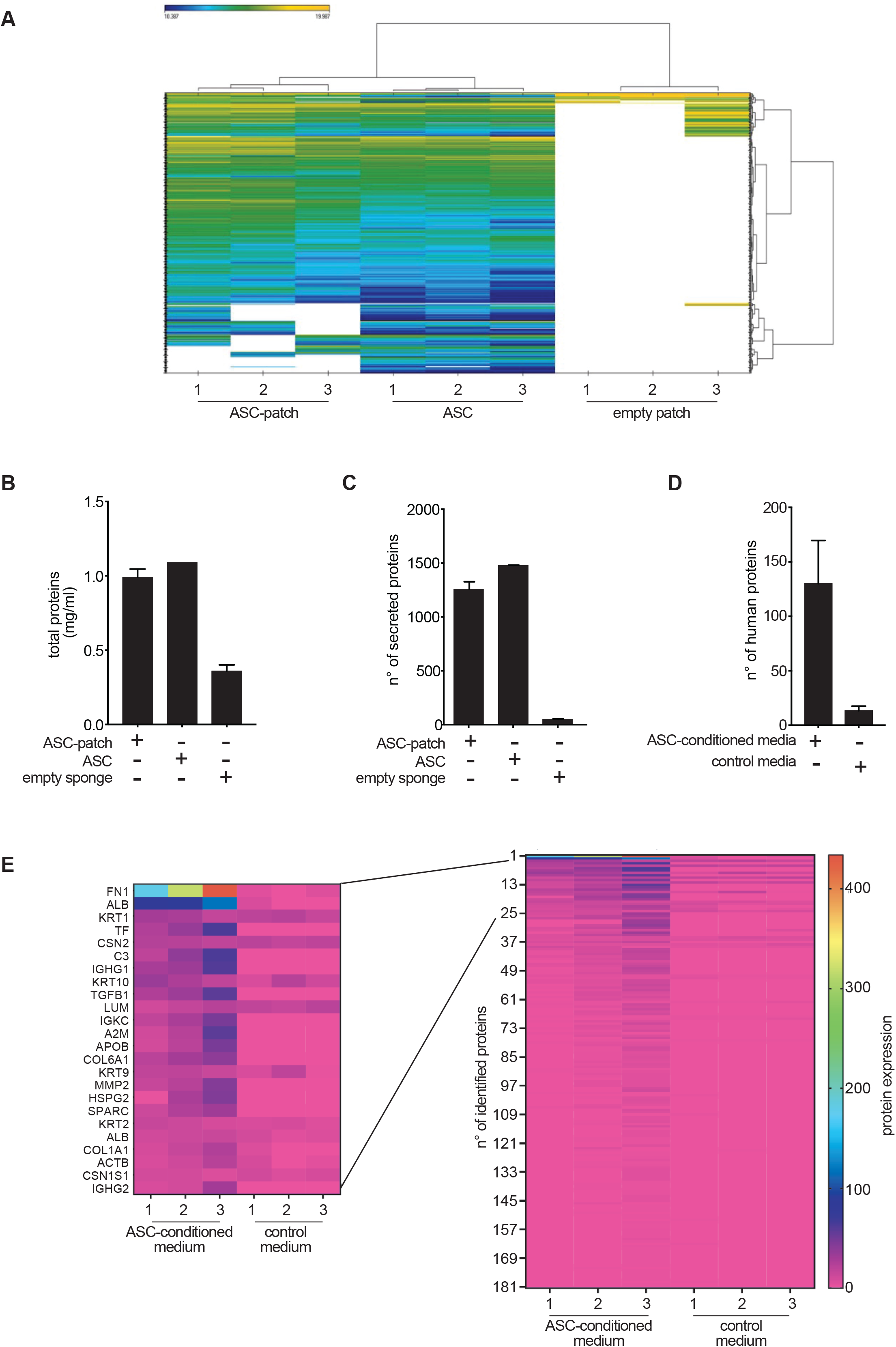
The ASC-patch concentrate the ASC secretome and gelatin-derived products. A-C) ASC-patches, ASC monolayer or empty gelatin sponge were incubated in a serum-free medium prior to a mass spectrometry analysis of the released proteome. ASC from three independent donors were analyzed. Hierarchical clustering analysis (A), total protein concentration (B) and total number of proteins identified (C) in each condition is shown. D, E) An empty gelatin sponge was pre-incubated with serum-free media preconditioned or not with ASC. After enzymatic digestion of the gelatin sponge, a proteomic analysis was performed by mass spectrometry. (C) Total number of proteins identified by mass spectrometry in each condition. (D) Heatmap showing the relative quantification of the proteins as shown in C, with a zoom on the top-25 most abundant. Overall in the figure, bars refer to mean+-SD.

We next assessed the ability of gelatin within the patch to adsorb proteins secreted by ASC. To this aim, three empty gelatin sponges were incubated with a serum-free medium or with a serum-free medium conditioned by ASC during 24h. Mass spectrometry analysis was performed on samples after complete dissolution of gelatin through collagenase treatment. Twelve proteins were identified in the samples deriving from gelatin sponges exposed to an empty serum-free medium. 181 proteins were instead detected in samples deriving from gelatin sponges exposed to the ASC-conditioned media (figure 3D), confirming that gelatin had adsorbed several factors produced by ASC. Figure 3E shows the relative abundancy of the 181 identified proteins in the two conditions, providing a zoom on the top-25. Interestingly, fibronectin-1 (FN1), a molecule important for the healing process [36] was the most abundant protein derived from ASC adsorbed on gelatin. The complete list of proteins is presented in supplementary table 4.

Together, these proteomic data indicate that the ASC-patch actively releases factors that are derived from both ASCs and gelatin. In addition, our results indicate that the patch function as reservoir of important healing factors, such as fibronectin, potentially promoting their slow and sustained release.

### The ASC-enriched patch promotes neo-angiogenesis, fibroblast survival and epithelialization

We next investigated the biological properties of the ASC-patch secretome on the growth of fibroblasts and keratinocytes, the main constituents of the dermis and epidermis, respectively. The survival of human fibroblasts cultured under poor nutritional condition, i.e without serum, was greatly enhanced by the addition of a conditioned media from the ASC-enriched patch (Figure 4A). Similarly, human keratinocytes could growth only when seeded above the ASC-enriched dressing, but not when applied to the empty gelatin scaffold, as confirmed by pan-keratin staining (Figure 4B, left and middle panels). Morphological analysis of histological sections highlighted the formation of a completed stratified epidermis. Of note, the basal layer of the neo-formed epidermis stained positive for the alpha chain of type XVII collagen (BP180, a structural component of hemidesmosomes), suggesting the formation of a correct basement membrane zone mediating adhesion of the epidermis to the ASC-enriched dressing (Figure 4B, right panel).

**Figure 4.**
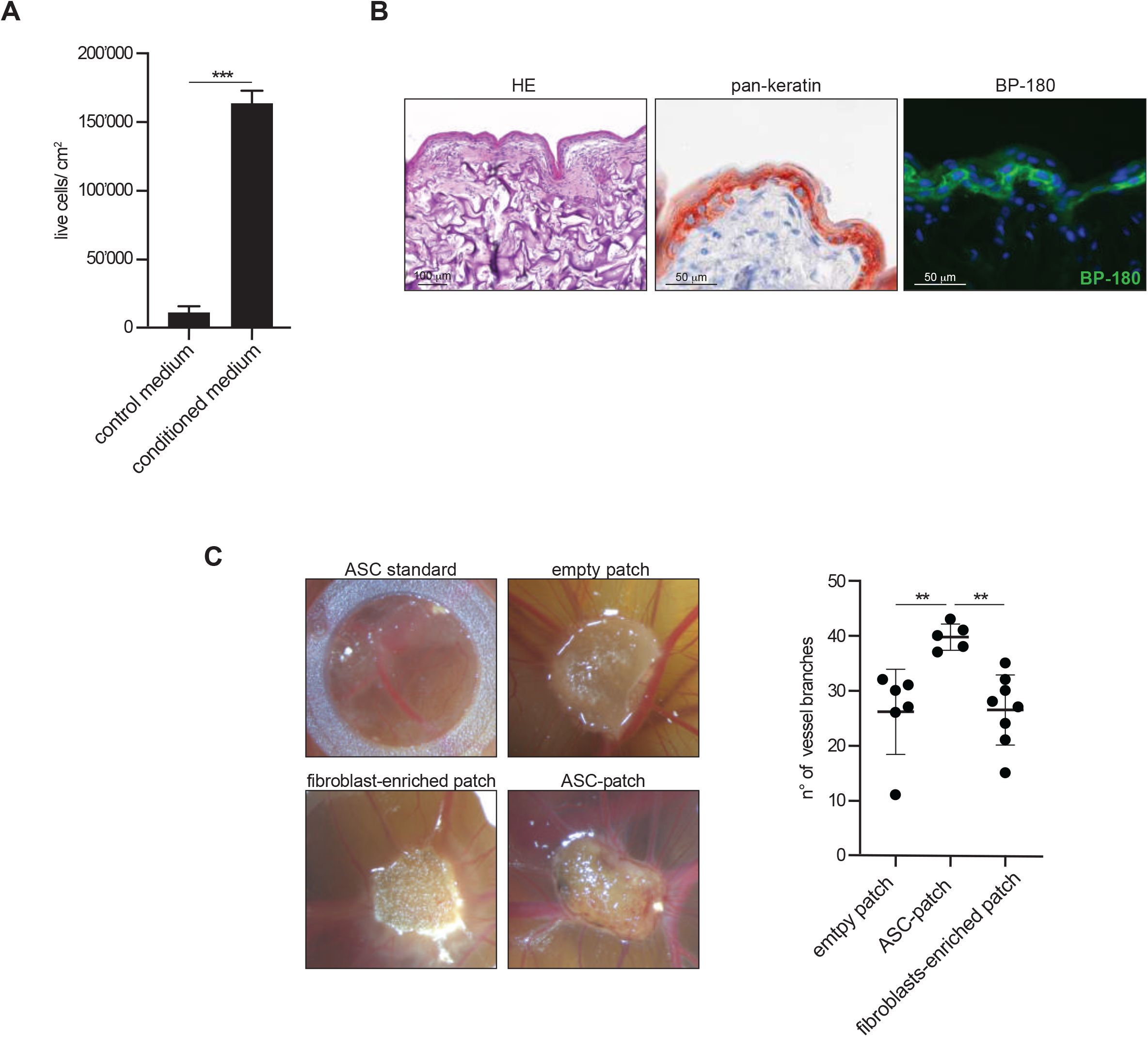
Figure 4. ASC-patches promotes neo-angiogenesis, fibroblast survival and epithelialization. A) Serum-free medium was incubated or not with ASC-patches generated from 3 different ischemic patient prior to its addition to human foreskin fibroblasts in monolayer. Cell viability was measured by trypan blue exclusion. B) Autologous primary keratinocytes were seeded on the ASC-patch for 15 days in air-liquid interphase. Shown is a representative hemalun/eosin coloration (left), pan-keratin immunohistochemistry (middle) and BP-180 immunofluorescence (right). C) ASC-patches, fibroblast-containing patches, ASCs in single cell suspension (via a silicon ring) or empty gelatin sponges were deposited on the chorioallantoic membrane of fertilized chick eggs. Representative images of neovascularization are shown for each condition (left). The number of new vessels connections were quantified using ImageJ (right). Overall in the figure: mean+-SD; nonparametric Mann Whitney t test: **= p<0.01, ***=p<0.001.

Neo-vascularization being a most critical process for wound healing, we next investigated the ability of the ASC-enriched dressing to promote angiogenesis *in vivo* by using the chicken choriollantoic membrane (CAM) model. To this aim, an ASC-patch was implanted on the CAM through a hole cut in chick eggshells, and its effect on the formation of new vessel branches was compared to that induced by an empty gelatin sponge, or a gelatin sponge containing human fibroblasts (Figure 4C). The ASC-patch was superior to all conditions tested, as a significant higher number of vessel branches sprouting from the dressing were observed (Figure 4C). Of note, new blood vessels could be visualized only into the ASC-enriched dressing by macroscopic analysis, but not the other conditions. A single cell suspension of ASC derived from standard culture was also applied by using a small circle of silicon to contain the cells and resulted in the absence of significant new vessel branches. The pro-angiogenic effect peculiar to the ASC-patch was probably due to synergistic mechanisms between the ASC and gelatin, since it was not merely the addition of the effect of its isolated constituents.

Together, we show here that the ASC-enriched dressing influences the biology of the most-important cutaneous cells and possesses angiogenic properties important to enhance the wound healing process.

### The ASC-enriched patch does not show tumorigenic activity and allows stabilization of ASC *in vivo*

We subsequently used immunodeficient Nude mice to investigate the tumorigenic potential of the ASC-patch. The experiments were conducted in accordance with the guidelines of the European pharmacopeia to study the tumorigenic potential of cell transplants. ASC-enriched dressings were subcutaneously transplanted in 10 Nu/Nu nude mice and compared with an equal number of mice transplanted with 5.10^6^ Hela tumor cells, this latter being the positive control for tumor development. At week five, 9 out of 10 mice that received Hela cells have developed palpable tumors, confirming the validity of the test. None of the mice receiving ASC-patch (5 sacrificed at week 5, the other 5 at week 12 according to European pharmacopeia guidelines) developed signs of palpable tumors in the transplant, in the peripheral tissues, nor in the brain, lungs, spleen, draining lymph nodes and kidneys as assessed by necropsy. Of note, all transplanted ASC-patch showed macroscopic vascularization at week 5, confirming its angiogenetic properties (figure 5A, left). Histological assessment of the transplanted ASC-patch revealed the presence of tissue-like cell-rich gelatin mesh (figure 5B). Culture of cells extracted upon enzymatic digestion of the transplanted ASC-patch (at week 5) had the morphology of ASC and expressed the Human Nuclear Antigen (HNA) (figure 5C). Flow cytometric analysis of this secondary ASC line confirmed that its ASC phenotype was conserved, as most cells expressed human CD73, CD90, CD105, CD44 and lacked the expression of CD45, HLA-DR and CD14 (figure 5D). Finally, the size of the transplanted ASC-patch was stable *in vivo* until week 3, and progressively decreased to almost be completely resorbed at week 12 (Figure 5A, right).

**Figure 5.**
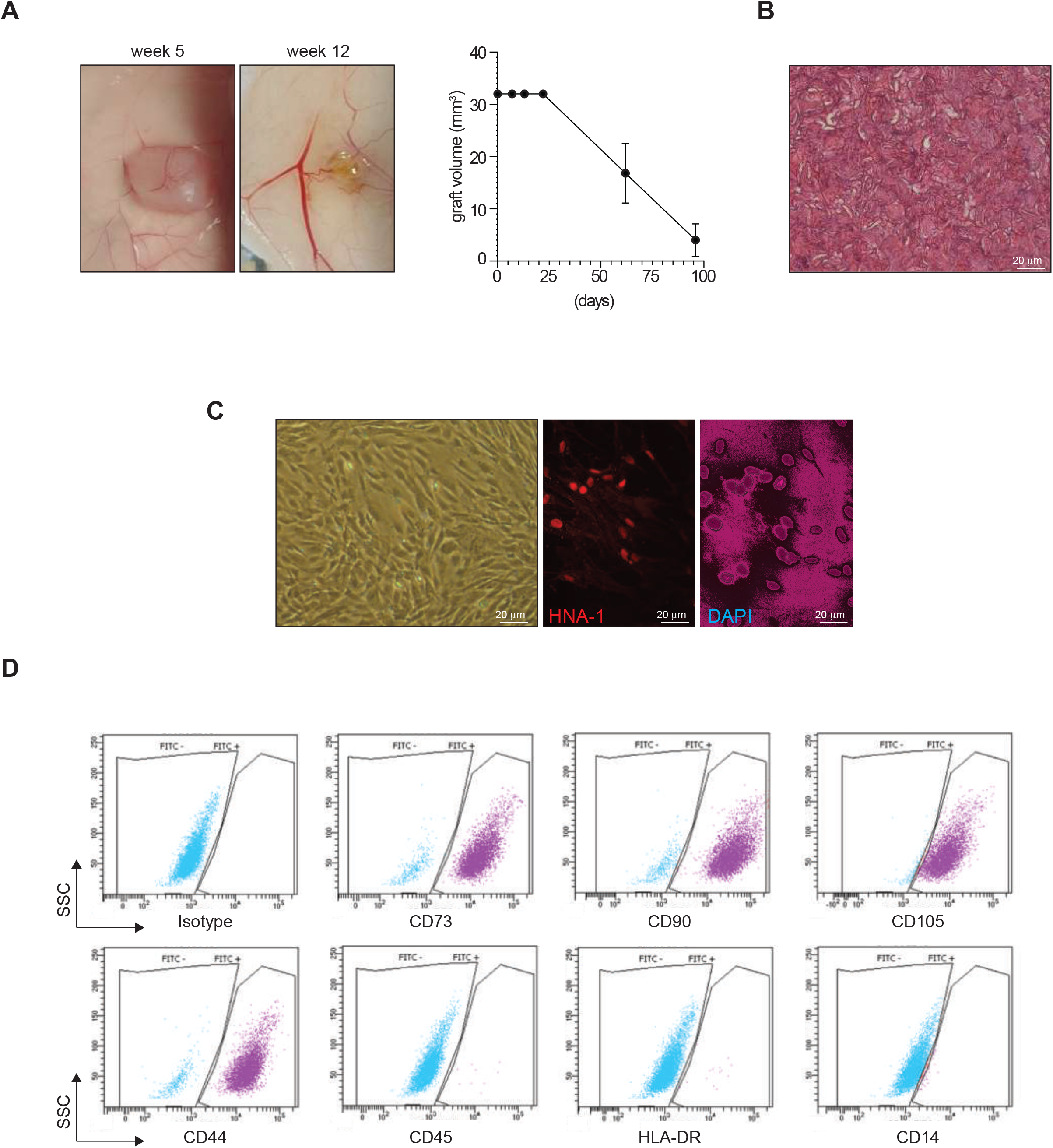
ASC-patches are stable in vivo after transplantation and do not show tumorigenicity. Ten Nu/Nu immunodeficient mice were transplanted subcutaneously with ASC-patches. Groups of 5 mice were sacrificed at weeks 5 and 12 for a macroscopic evaluation of transplants. A) representative images of transplanted ASC-patches at week 5 and 12 (left) and quantification of graft volume over time (right). B) Hemalun/eosin coloration of a transplanted ASC-patch at week 5. C-D) Cells were extracted after dissociation of grafted ASC-patches at week 5. C) Representative images of the morphology (left), and Human Nuclear Antigen (HNA)/DAPI immunofluorescence (middle and right) of the cells extracted from the transplanted ASC-patches. D) representative flow cytometry analysis of the extracted cell line for the indicated ASC markers (positive: CD73, CD90, CD105; negative: CD45, HLA-DR, CD14).

Together, these *in vivo* experiments indicated that the ASC-patch lacks any tumorigenic activity and allow the stabilization of the ASC for at least 5 weeks, while the whole product is resorbed within 12 weeks in an immunodeficient setting. The ASC-patch has thus essential pre-clinical safety requirements for further development in humans.

### The ASC-patch boosts the healing process and induces early neo-angiogenesis in a rat model of ischemic wound

Next, we investigated the *in vivo* efficacy of the ASC-patch in a pre-clinical rat model of ischemic wound [25]. Wounds were created on the dorsal part of the hind paws of Wistar rats by removing a full□thickness skin area. To create paw ischemia, 1 cm of the femoral artery was surgically removed prior to the wound creation. In this experiments, ASC-patches were prepared from syngeneic rat ASC to avoid rejection in immunocompetent rats (rASC-patch). Identity of rat ASC was confirmed by flow cytometry (data not shown). Like in experiments using human cells, rat ASC-patches were characterized by an enhanced regenerative transcriptome when compared to rASC grown in standard monolayer conditions as assessed by microarray (data not shown). Of note, rats treated with rat ASC-patches healed faster than rats treated with empty gelatin patches or standard silicone/polyurethane dressings confirming the efficacy of the ASC-enriched dressing *in vivo* (Figure 6A and B). A granulation tissue was macroscopically visible from day 9 in the ASC-patch treated group, while tendons were still exposed in the control groups (Figure 6A). All rats treated with the ASC-enriched dressings reached a complete wound closure before day 17, compared to 67% of rats treated with the empty gelatin sponge (Figure 6B and data not shown). In line with the previously observed effects on angiogenesis in the CAM model, staining for the endothelial marker CD31 revealed an increased vascularization in the healing tissue of rats treated with ASC-patch as compared to controls (Figure 6C). Vessels, which were increased in numbers, had a generally higher diameter and appeared structured in a denser network. Quantification of the total vessel area by image analysis confirmed the superiority of ASC-patch rather than empty gelatin sponge and standard dressing (figure 6D).

**Figure 6.**
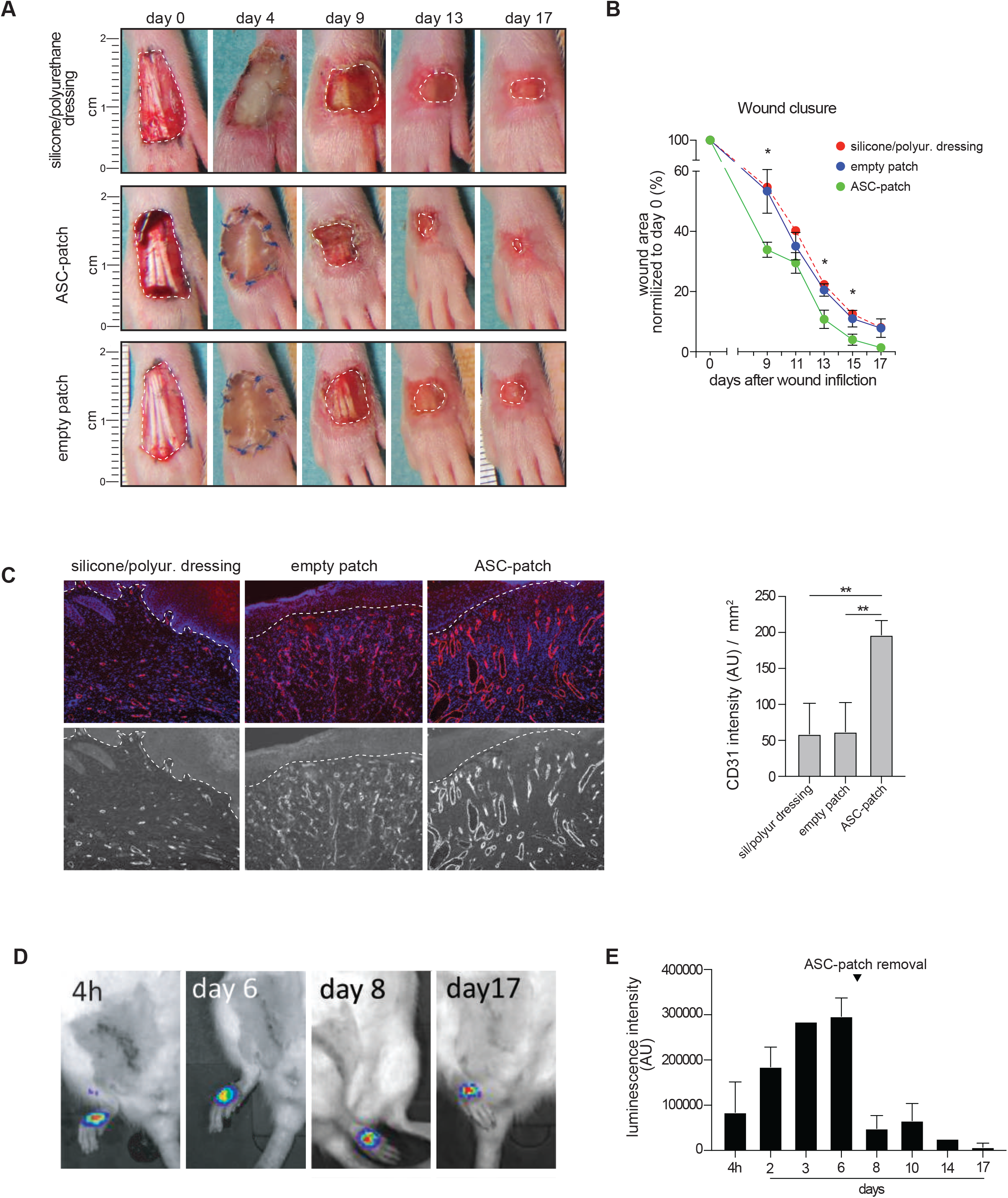
ASC-patches close ischemic wounds faster than acellular gelatin sponge and silicon/polyurethane dressings, with an increased vascularization of the granulation tissue. ASC-patches, empty gelatin sponge or a standard dressing of silicone/polyurethane were sutured on a full thickness wound on the ischemic foot of Wistar rats. Treatment was kept in place for the initial 7 days. A) Representative macroscopic images are shown at different time points. B) Quantification of the wound area, as normalized to day 0. Shown are cumulative data from 6 ASC-patches and 5 empty patches. A representative silicone/polyurethan dressing is shown as control. C) representative CD31 (endothelial cell marker) immunofluorescent staining of the granulation tissue at day 7 (left). Quantification of the CD31 intensity/ mm^2^ (right). D) Representative live imaging of rASC _EFS FLuc_ after intraperitoneal injection of D-Luciferin. Images are taken one hour after the substrate injection. E) Quantification of the luminescence emitted in the wound zone over 17 days post-treatment. Cumulative data from 3 rats is shown. Mean+_SD; non-parametric Mann Whitney t test: **= p<0.01; *p<0.05. AU: Arbitrary Units

Finally, the survival of ASCs within the patch was analyzed *in vivo*. Rat ASCs stably transduced with firefly luciferase (FLuc) under the control of the ubiquitous promoter EFS (rASC _EFS FLuc_) were used to generate rar ASC-patches (FLuc-rASC-Patch). Intraperitoneal injection of D-luciferin allowed the monitoring of ASC survival in vivo by using the live imaging system IVIS Spectrum (Perkin Elmer). Luminescence in the FLuc-ASC-Patch was detected until removal of the treatment at day 7, confirming survival of ASC *in vivo* (figure 6D and E). Luminescence was maintained after treatment removal until day 17, indicating that some ASC engrafted within the healing wound. Additional quantifications during the entire period of treatment indicated a fast increase of the luminescence until day 6, an observation in favor of rapid exchanges of the patch with the biological fluids and the increased neovascularization previously observed (figure 6E).

Together, we provide evidence that the ASCs within the ASC-patch are stable and viable in vivo for at least 17 days. Of note, the ASC-patch accelerates wound healing in a rat model of ischemic wounds, resulting in increased neo-vascularization and emergence of a granulation tissue earlier in the healing process.

## DISCUSSION

In this study, we demonstrate the *in vitro* regeneration potential and pre-clinical efficacy/safety of a cellular patch composed of ASCs embedded in a bovine crosslinked-gelatin sponge. Extensive molecular and biochemical characterization revealed that induction of angiogenesis, in the absence of tumorigenesis, is one of the most important mechanisms of action of this approach. The patch formulation represents an optimal non-invasive delivery route that maximizes the local effects of mesenchymal stem cells within the wound. This study validates the patch approach in the pre-clinical setting and paves the way for its use in first-in-human studies.

The ASC-patch studied here essentially relies on the combined interaction of adipose-derived stem cells with gelatin. ASCs are known for their ability to enhance the healing process by secretion of soluble healing factors and extracellular vesicles [37-40]. Here we demonstrate that ASCs not only increase their regenerative potential when they grow within the patch, but are also locally concentrated. Of note, gelatin and collagens influence the cell function directly [29]. ASCs exposed to gelatin microcryogels have been shown to increase their gene expression of VEGF, HGF, FGF, and PDGF [41]. Similarly, gelatin-exposed fibroblasts increased their proliferation, differentiation towards myofibroblasts, and cytokeratin secretion [42]. In addition, the patch allows for the formation of a three-dimensional tissue organization that enables the optimization of cell-cell interactions and paracrine events [43]. Finally, the condition used for culturing the ASC-patch, i.e. air/liquid interface, promotes air gas exchanges and oxygen supply, which are a key factor influencing ASC functions [44].

The gelatin sponge itself in the ASC-patch may promote the healing process independently of ASCs. Gelatin is haemostatic, acts as a chemotactic agent [29], promotes the formation of a granulation tissue [28] and absorbs exudates present in the wound bed, creating a microenvironment favouring the the inflammatory phase of wound healing [27]. In line with this data, gelatin was shown to directly enhance wound closure [29], particularly by increasing angiogenesis, keratinocyte and fibroblast proliferation / migration and myofibroblast differentiation [30,42,45]. Besides the possible direct action of gelatin within the ASC-patch, we have shown that the gelatin patch slowly dissolves in aqueous culture conditions, probably under the influence of ASCs, releasing numerous collagens and cytokeratins. As gelatin, collagens and cytokeratins promote wound healing through enhancement of proliferation and migration of fibroblasts and keratinocytes, acting also on IL-1, VEGF and TNF-α secretion [46-52].

Previous studies have shown the possibility to include MSCs in acellular scaffolds *in vitro* [53] and confirmed their clinical potential for cartilage repair [54], annulus defect [55] or spinal cord injury [56]. A dermic substitute of collagen or atelocollagen loaded with MSC, was also tested in animals [57] or patients [58], showing an enhancement of wound closure without any toxicity. In this work, we have used gelatin and considered it ideal to generate the ASC-patch for wounds. First, gelatin is a biodegradable and biocompatible scaffold that is already widely used in clinical settings. In fact, gelatin sponges or gelatin powders are used for bleeding control in surgical procedures and are well known to induce no antigenicity or toxicity in cells and patients. Gelatin, which consists of more than 90% protein, mineral salts, and water [59], is produced by hydrolysis of the triple-helix structure of animal collagens through heat or enzymatic denaturation, producing random coiled domains. Compared with collagens or atelocollagens (a collagen derivative without telopeptides at the end of collagen fibers), gelatin lost the typical triple-helix organization of the fibers but retained a molecular composition very similar to that of collagen, thus maintaining the important binding structures for cellular attachment. Unlike collagen, gelatin is highly soluble in water, an essential feature for cell therapy, and is relatively inexpensive.

To create the ASC-patch, we adopted a sponge formulation of gelatin. The porosity of the sponge is indeed particularly attractive to allow ASC growth within the pores, simultaneously permitting the biological fluids to circulate, as well as the colonization by host cells and vessels. In line with this, we have observed a very rapid irrigation of the sponge when applied in vivo on ischemic non-exudative wounds and a colonisation of the patch by host cells. The malleability and physical stability of gelatin sponges allows their spatial adaptation to the wound bed, an important prerequisite for the clinical use, regarding the high diversity of the shape and depth of chronic wounds in humans.

The primary purpose of this study was to demonstrate the efficacy and safety of a novel formulation of ASC-patch for the treatment of chronic refractory ulcers. Our observations suggest that ASC-patch promotes wound closure thought increased angiogenesis, as well as fibroblast survival and epithelialization. The mechanisms of action of the ASC-patch are based on multiple effects, including the direct effect of gelatin, gelatin-derived products, local concentration of ASCs, and a controlled release of the ASC secretome locally into the wound. The ASC-patch lacks tumorigenicity, as confirmed by a European Pharmacopoeia-recommended test, and has no adverse effects or pro-inflammatory effects in animals.

The ASC-patch formulation shown in this study represents an attractive non-invasive delivery route which maximizes the local efficacy of MSC therapy for chronic wound care, as shown *in vitro* and in animal models. First-in-human studies are planned in the near future to clinically validate the ASC-patch solution as a simple and effective treatment improving patient compliance and physician acceptance.

## LIST OF ABBREVIATIONS

ASC: Adipose-derived Stem Cells
CAM: chicken choriollantoic membrane
FLuc: Firefly luciferase;
GMP: Good-Manufacturing Practice
MMP: Matrix Metallo-Proteinases
MSC: Mesenchymal Stromal Cells
PCA: Principal Component Analysis

## Supplementary Figure Legends

**Supplementary figure 1.**
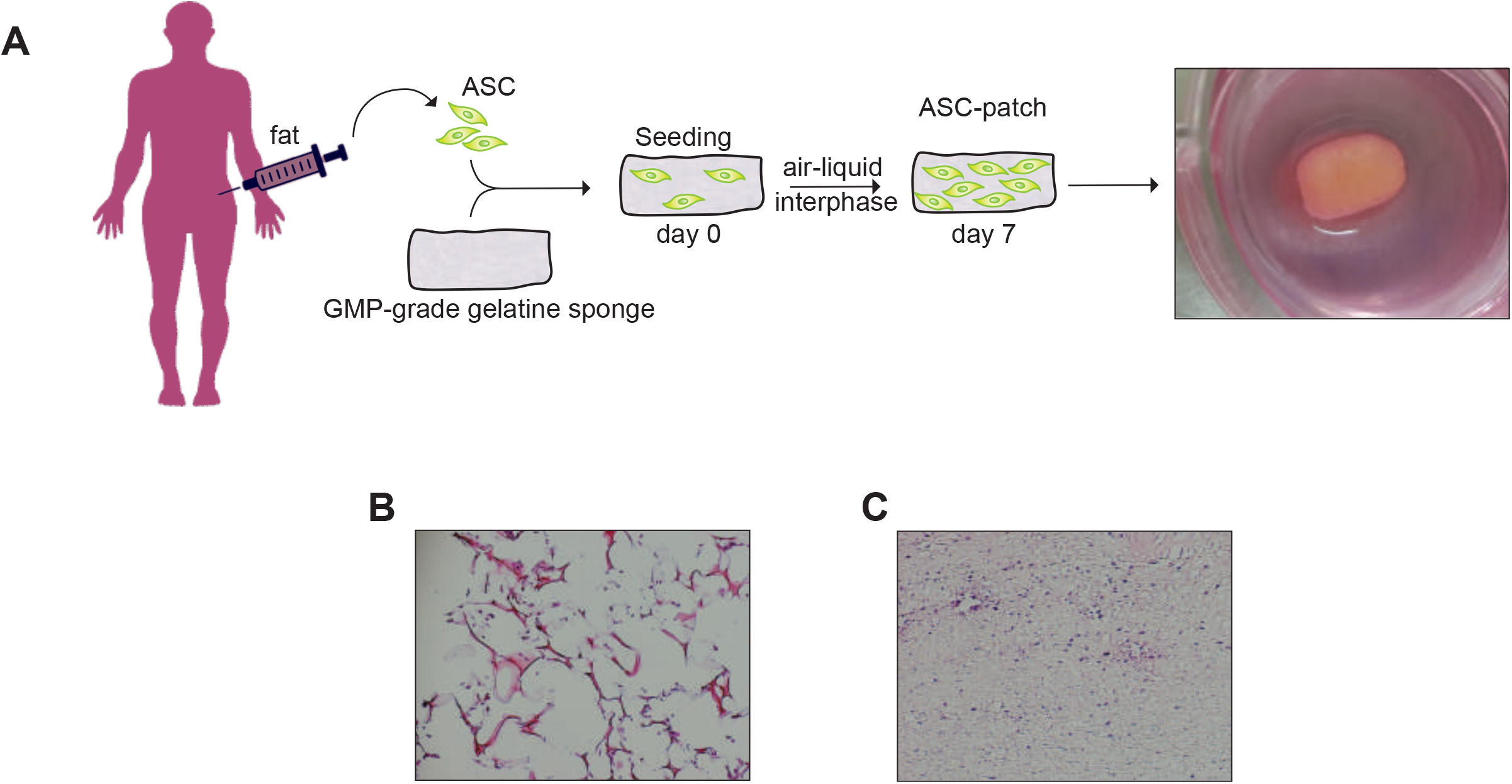
Generation of the ASC-patch. A) ASC derived from the abdominal fat and introduced within gelatin sponges and cultured during 7 days in air/liquid interface conditions to generate ASC-pacth. B) Hemalum/eosin coloration of an ASC-patch at day 0, immediately after ASC seeding. C) Hemalum/eosin coloration of an ASC/collagen gel composite 24h after ASC inclusion. B,C) images representative of 3 experiments.

**Supplementary figure 2.**
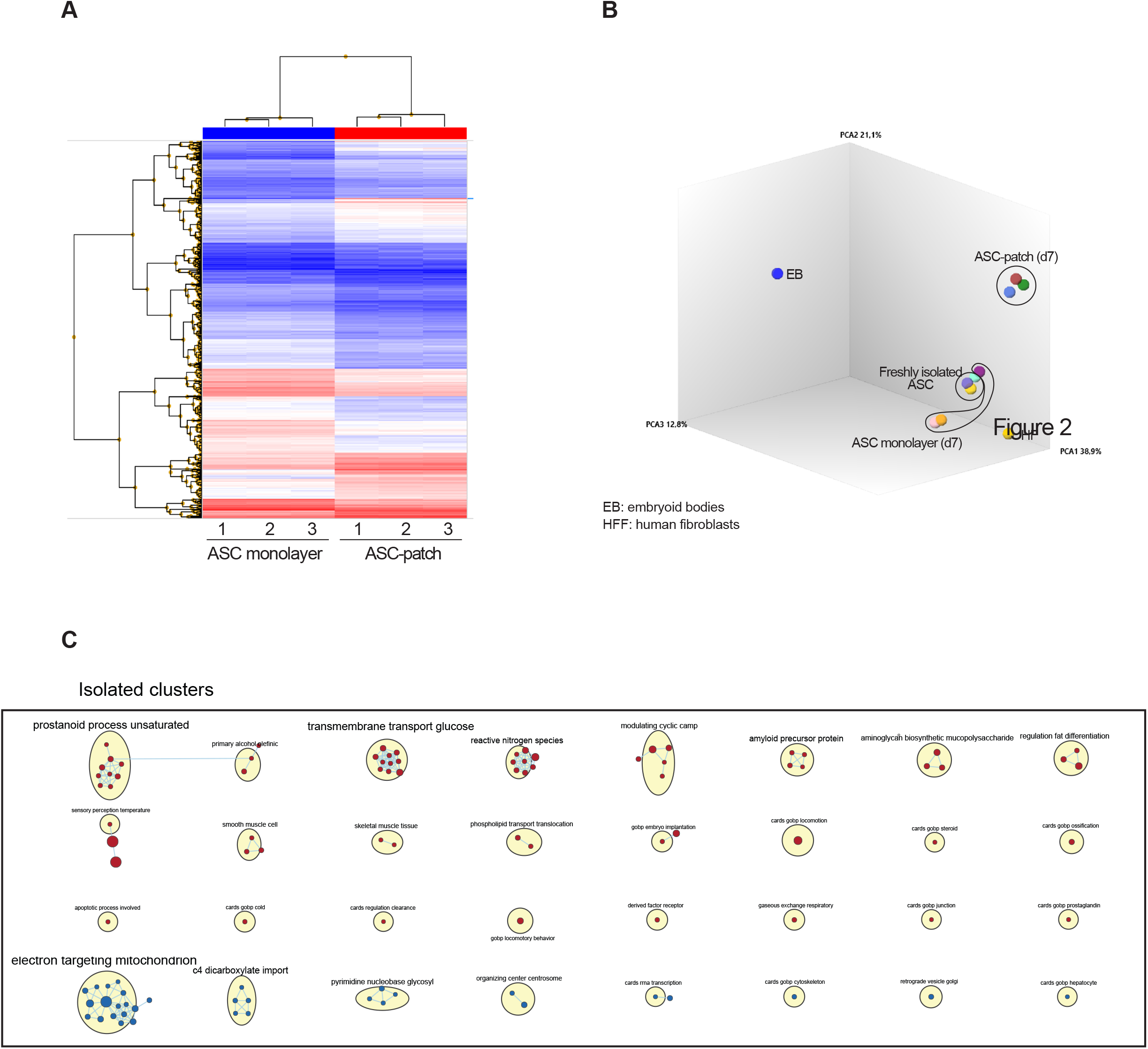
ASC in gelatin sponge regulate their regenerative transcriptome. The transcriptome of ASC-patches generated from 3 ischemic patients were assessed by microarray and compared to ASC from the same donors grown in monolayer. A) Hierarchical clustering of the two different conditions. B) Principal Component Analysis (PCA) of the microarray data of ASC-enriched patch (at day 7), ASC grown in monolayer (at day 7), freshly isolated ASC, embryonic bodies (EB) and human fibroblasts (HF). C) Isolated and less represented clusters from analysis shown in Figure 2.

**Supplementary figure 3.**
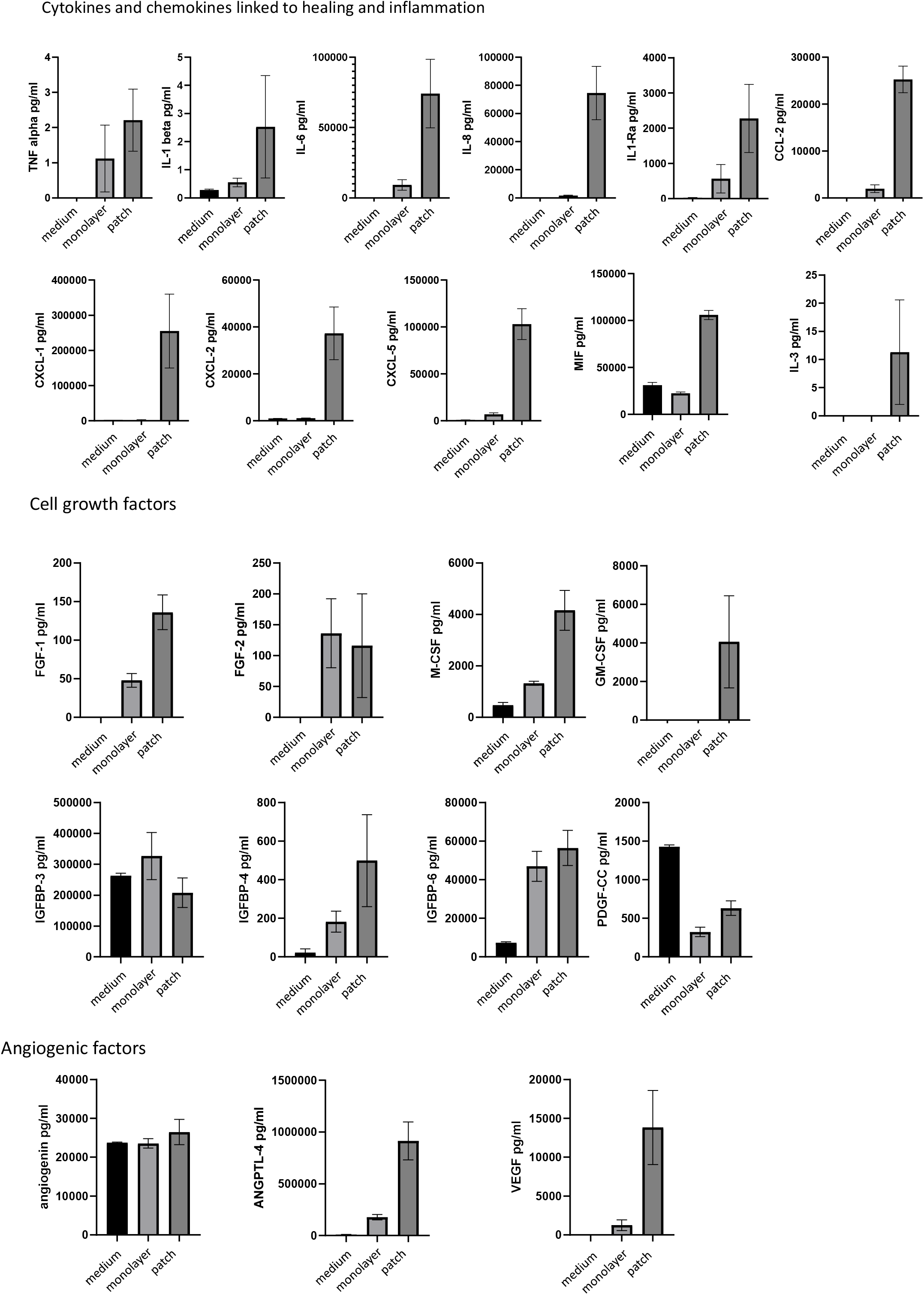
Healing factors concentrated in the ASC-patch supernatants. Quantification of the protein level of the indicated healing factors by bio-array in the supernatants of ASC-patches and ASC grown in monolayer.

**Supplementary figure 4.**
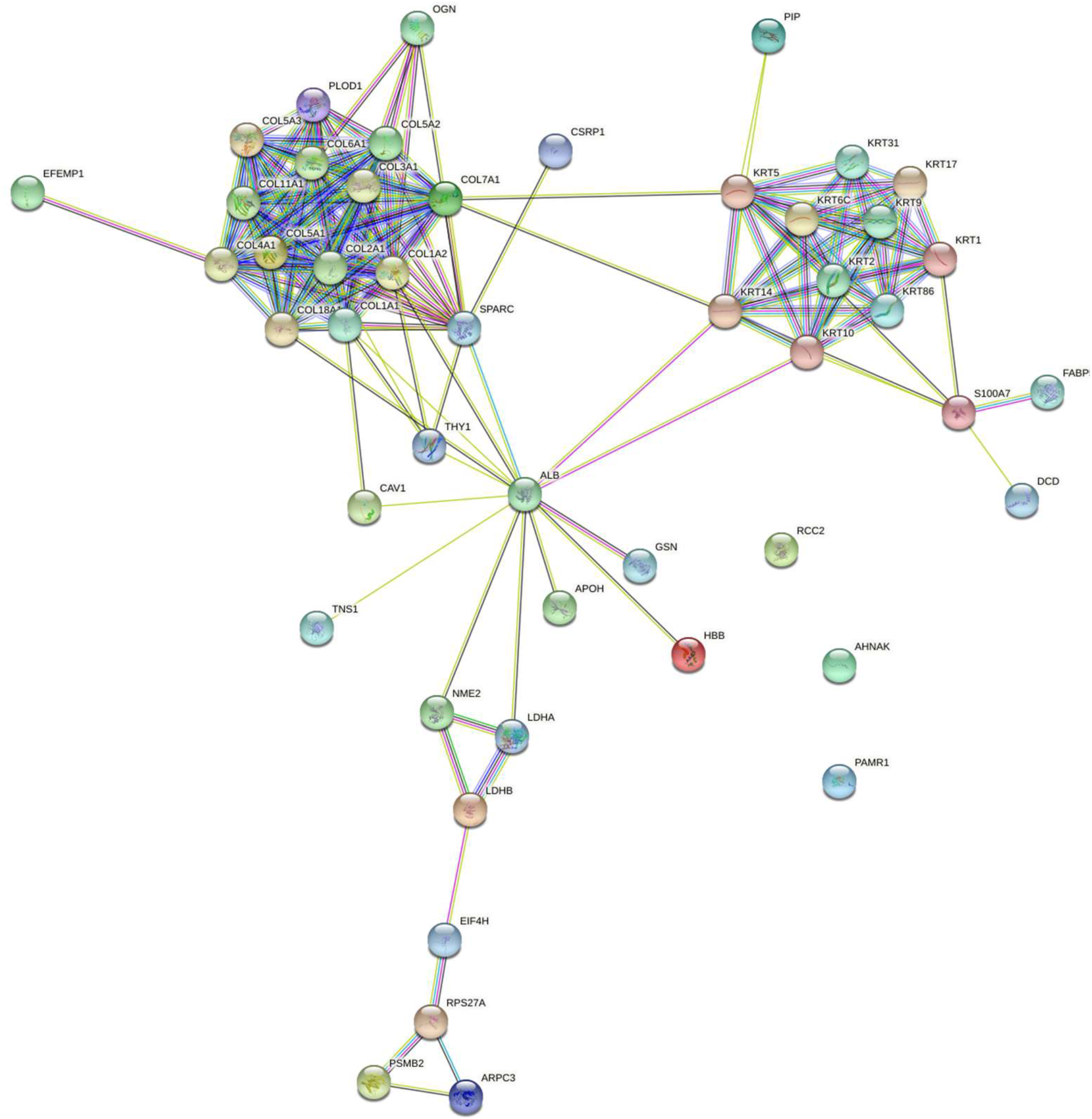
The acellular gelatin sponge by itself releases collagens and keratins. Empty gelatin sponges were incubated for 24h in a serum-free medium, prior to a mass spectrometry analysis of the released proteome. Clustered protein according stringDB are shown. The experiment was performed in triplicates.

**Supplementary Table 1**

Raw data of microarray comparing the ASC-patch condition and the ASCs grown in monolayer.

**Supplementary Table 2**

List of the 191 gene transcripts corresponding to the most implicated factors in the wound regeneration analyzed in more details.

**Supplementary Table 3**

List of proteins found in the supernatants of ASC-patches and ASCs grown in monolayer.

**Supplementary Table 4**

List of the 181 proteins detected in samples deriving from gelatin sponges exposed to the ASC-conditioned media.

## Acknowledgments

The authors would like to thank Barbara Russo (Dermatology, University Hospitals of Geneva, Switzerland) for its help in microarray data analysis. This work was supported by “Fondation pour la recherche sur le diabète”, Innosuisse (grant n° 43333.1), the University of Geneva (INNOGAP), “Fondation Louis Jeantet–Fondation privée des Hopitaux Universitaires de Genève (STARTER). Data available on request from the authors.

