## supplementary figures for "Efficacy and safety of a patch containing adipose-derived stem cells for skin wound healing – results form a comprehensive pre-clinical evaluation program"

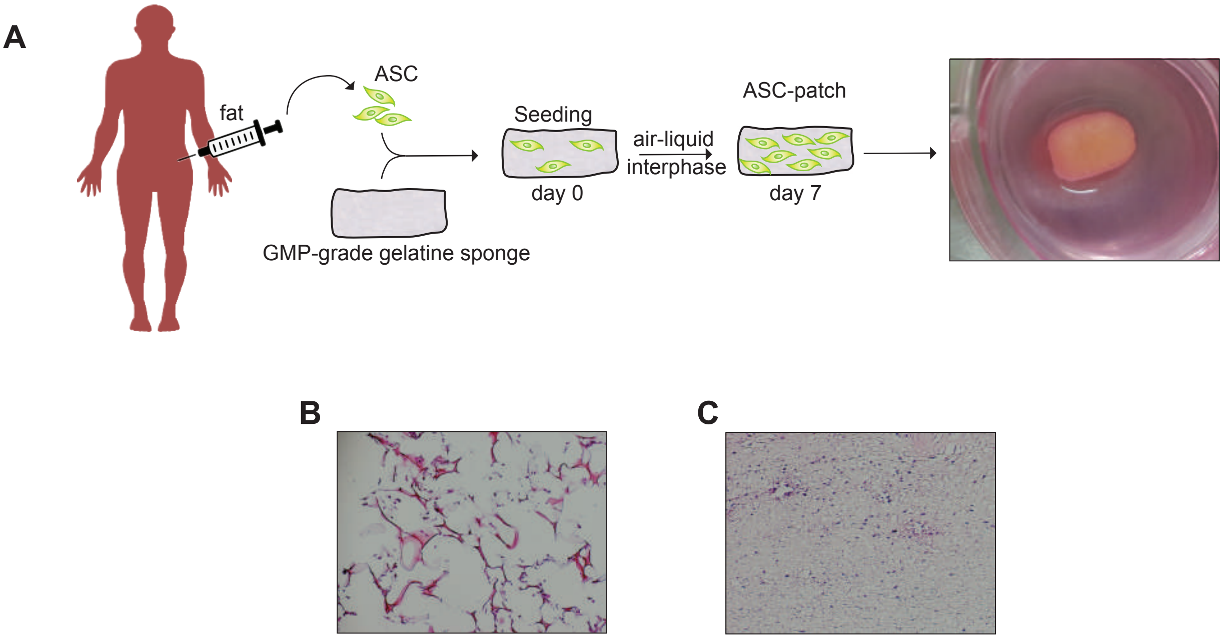


Supplementary Figure 1

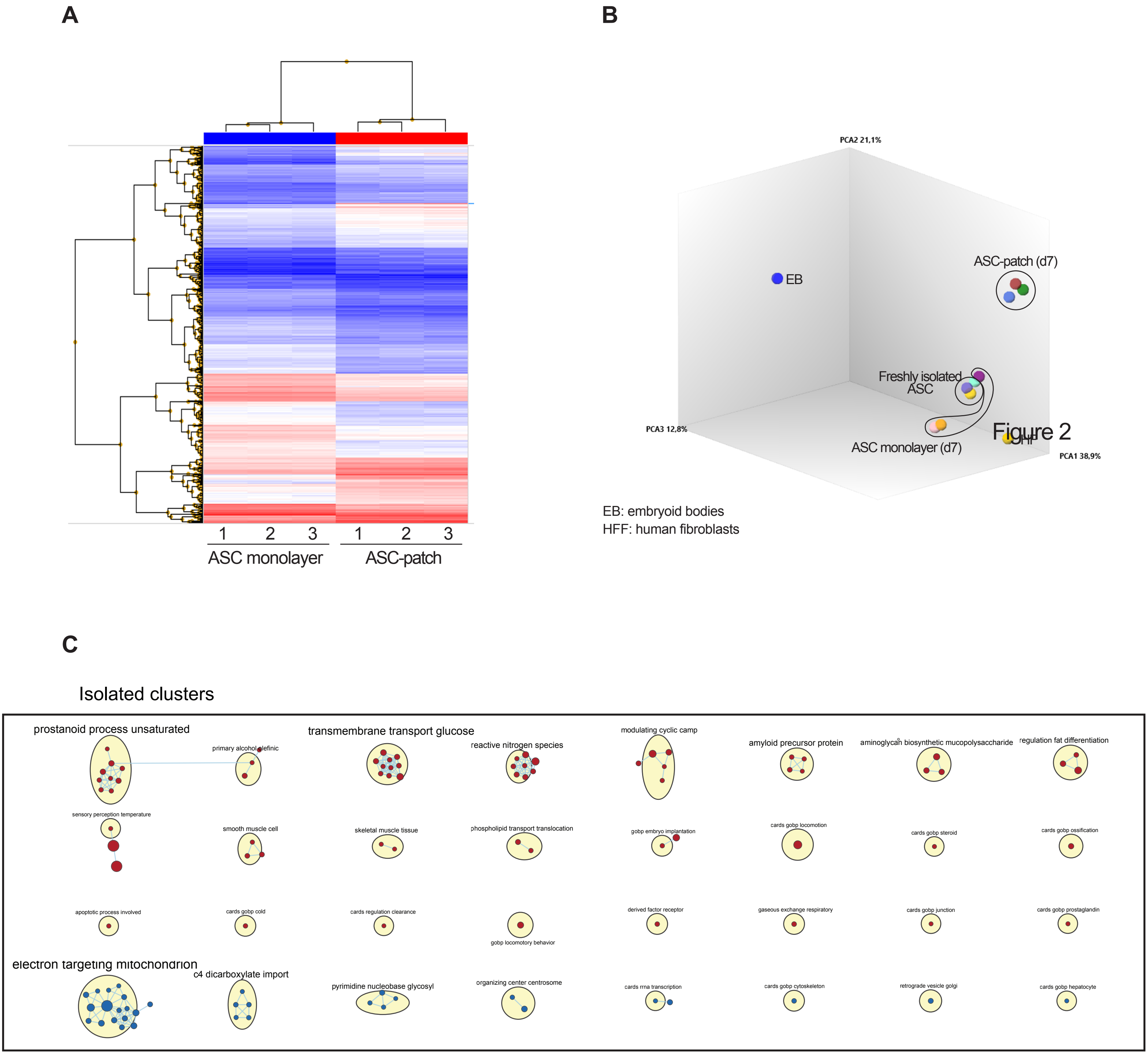


Supplementary Figure 2

Cytokines and chemokines linked to healing and inflammation

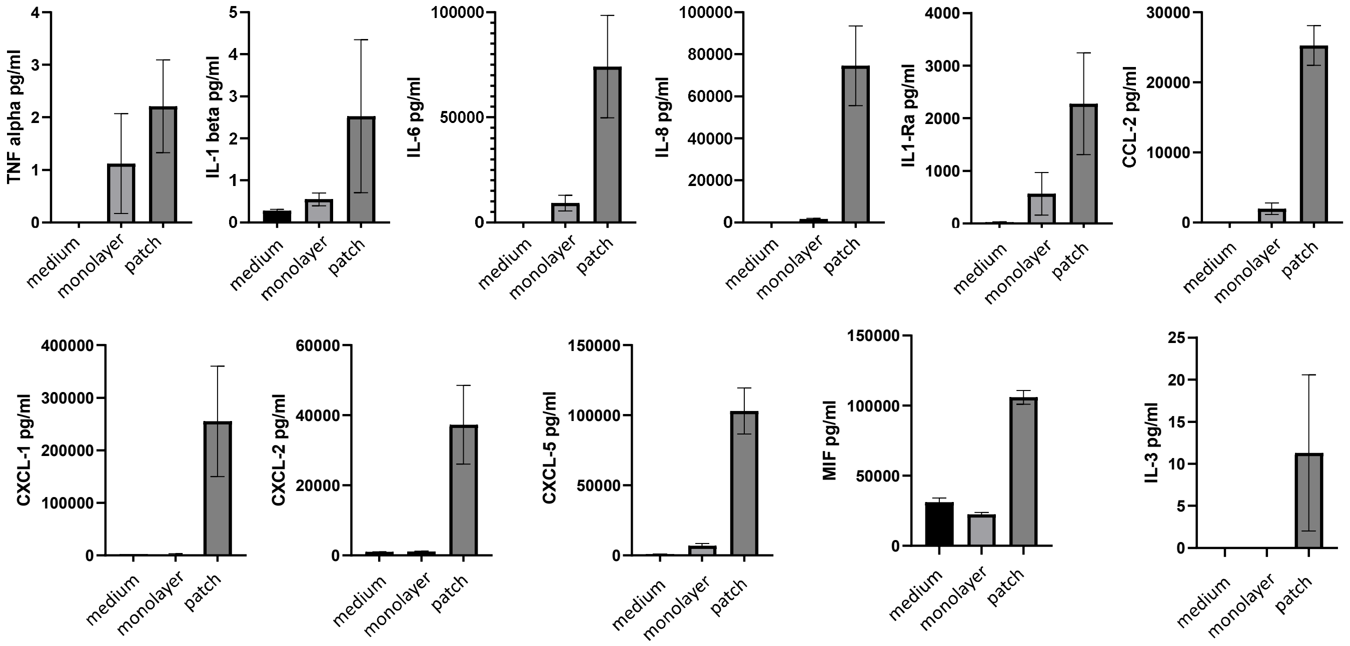


Cell growth factors

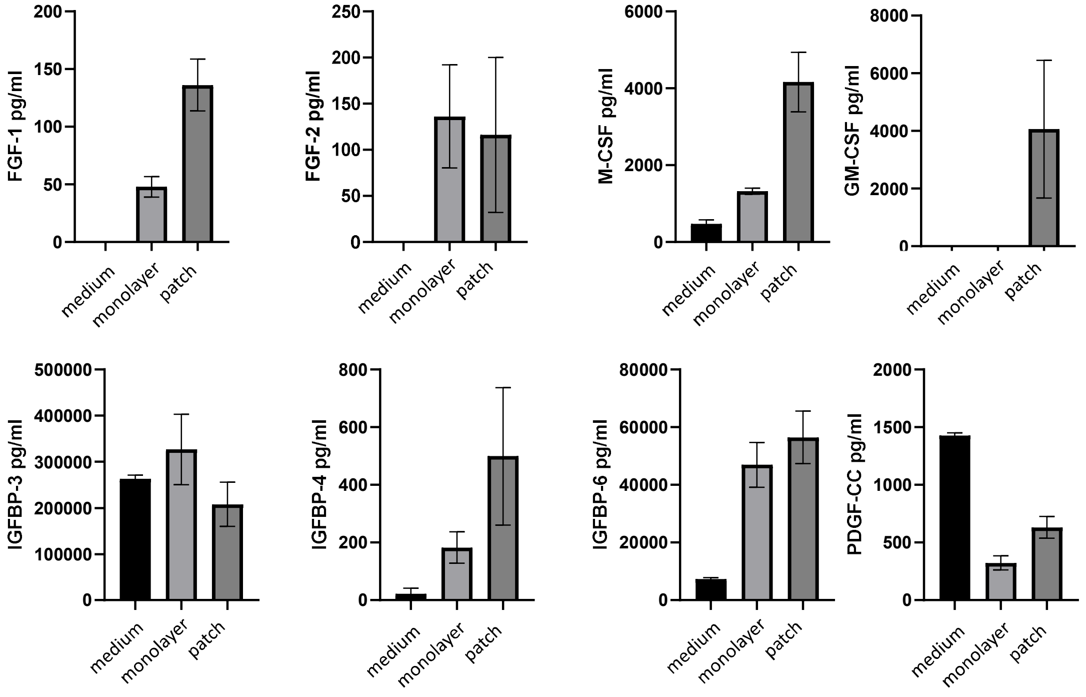


Angiogenic factors

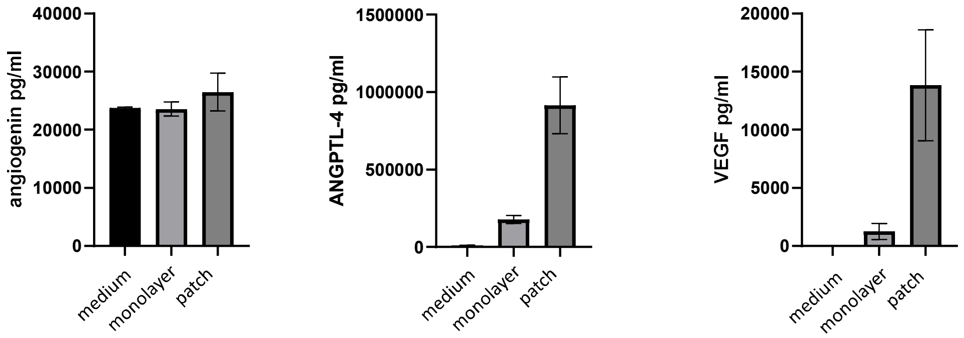


Supplementary figure 3

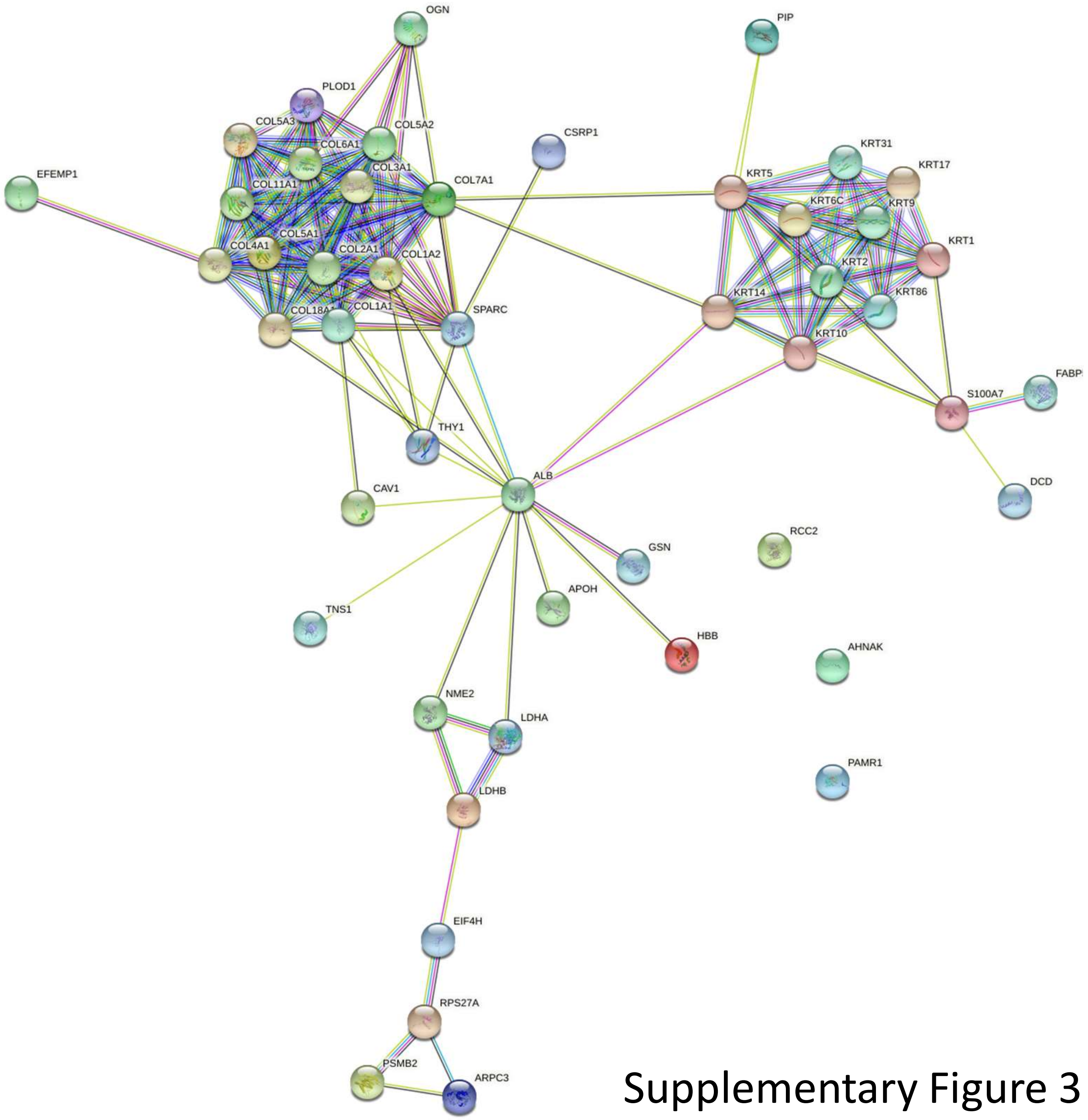


Supplementary Figure 3

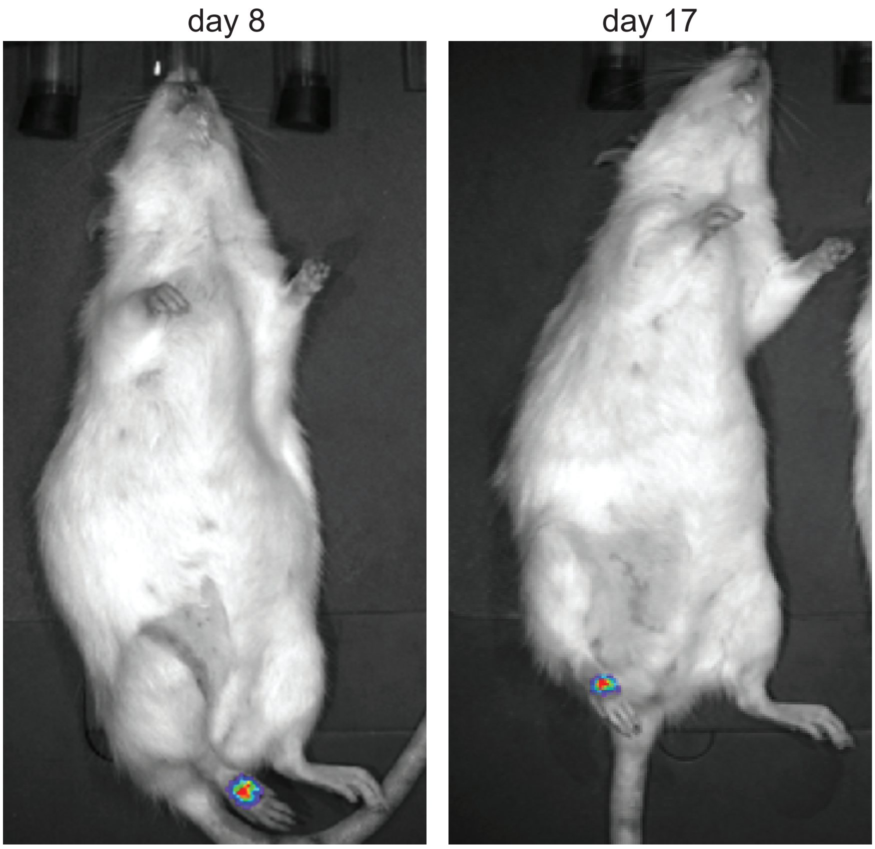


Supplementary Figure 5
